# Highlighting the strengths of new wearable fNIRS, precision accuracy microgenetic research designs, and new NeuroDOT processing algorithm to measure change over time in clinical populations for Speech-Language and Hearing Researchers

**DOI:** 10.64898/2026.06.09.725667

**Authors:** Sarah Crow, Ari Segel, Emma Speh, Paulina Skolasinska, Adam Eggebrecht, Julia L. Evans

## Abstract

Fundamental to understanding the process of intervention among individuals with communication disorders is the ability to document behavioral changes over time. This technical report demonstrates the clinical applicability of wearable fNIRS systems and NeuroDOT processing pipelines for examining within-person cortical dynamics of learning. Using a microgenetic research design and a dense sampling approach, we examined changes in the prefrontal cortical hemodynamic response in an adult female participant who completed the same spoken sentence repetition and auditory fixation tasks across eight sessions. In addition to participant accuracy, hemodynamic data were collected with a continuous-wave, multi-channel fNIRS system (NIRSport2) using a prefrontal 20-channel optode montage. Data were processed using NeuroDOT (https://www.nitrc.org/projects/neurodot) (Eggebrecht & Culver, 2019) to: (i) standardize signal quality across the sessions to quantify motion levels and to ensure standardized brain map specificity, (ii) to examine both channel space fluctuations in the hemodynamic response and map changes in cortical activation patterns over the sessions. The signal quality met the predefined criteria for only the first five sessions. Participant’s repetition accuracy did not improve over the five sessions. Channel-wise analysis revealed that HbO concentration differs significantly over right and left hemisphere channels over the course of the five sessions for the Sentence Repetition task, but not for the Auditory Fixation condition. Brain maps revealed qualitative differences in the pattern of prefrontal cortical activation across the five sessions. To fully document the mechanisms that drive changes in behavior, understanding the brain-behavior link is critical to identifying and discriminating between typical and atypical learning.

## 1. Introduction

The purpose of this technical report is to demonstrate how wearable functional near-infrared spectroscopy (fNIRS) optical imaging methods, when combined with recently developed optical brain imaging data processing algorithms and a dense-sampling microgenetic research design, can provide unique insights into the dynamics of learning among individuals with communication disorders. Effective clinical interventions require an understanding not only of how the learning trajectories among individuals with communication disorders differ from those of typical individuals, but also how they differ within a clinical population. This necessitates methods that enable the real-time examination of within-individual learning trajectories. While cross-sectional and/or longitudinal research designs excel in describing differences in the language abilities among atypical and typical learners at different ages or stages in the progression from novice to expert (e.g., Biró et al., 2025; Pennington, et al., 2021; Kolb & Kolb, 2018; Oogarah-Pratap, et al., 2025), they are not well suited to identifying the mechanism(s) that underlie the individual learning trajectories of atypical and typical learners.

The focus of microgenetic research designs is on *how* an individual learns and if skill X is needed before an individual can learn Y (Siegler & Crowley, 1991). Fundamental to the microgenetic approach is characterizing the dynamic balance between learning and stable performance (e.g., competence). For example, childhood is characterized predominantly by “learning,” whereas adulthood is characterized by a shift into more stable performance state(s). At the microscopic level, understanding the mechanism(s) that underlie this shift from learning to stable competence in real-time during the course of therapy is critical to the development of effective interventions. Notably, cross-sectional and/or longitudinal research designs assume that “learning” is characterized by a steady, linear progression from less to more advanced states of competence. However, a microgenetic approach not only assumes that learning is characterized by the appearance of regressions and progressions, but that there are temporary transitional states that will be present only briefly and that can only be identified through multiple levels of observation (Seigler, 2006). Finally, to understand how the learning trajectories of atypical learners differ, the microgenetic approach assumes that observations need to start *prior* to the emergence of the new skill or behavior and *continue* through the entire period of rapidly changing competence.

In microgenetic designs, the frequency of data collection must be proportional to the rate of change of the behavior that is being studied. Siegler introduced the term “dense” sampling as a core property of this high-density observational approach. Thus, capturing this progression requires a “dense” sampling approach with sufficient temporal resolution (e.g., timing of the observations) to capture regressions, progressions, and temporary transitional states in real time. D’Souza and Karmiloff-Smith (2017) argue further that to understand how the “learning” trajectories of individuals with communication disorders differ from those of typical individuals also requires understanding how the material to be learned uniquely interacts with the individual learner. They also argue that qualitative differences between atypical and typical learners can only be identified if observations begin at the earliest point in the divergence of the learning trajectories and follow their trajectories with dense observation paradigms, such as those characteristic of microgenetic approaches. They also argue that identifying these early divergent time points requires observations across multiple levels of description (e.g., genetic, neural, behavioral, environmental). Although the goals differ, the field of neuroimaging has begun to take a similar approach. For example, precision neuroscience brain imaging is an emerging approach that uses dense data collection (typically MRI) to acquire detailed, individual-specific brain maps to understand the unique characteristics of brain structure and/or function (Demeter & Green, 2025). Similarly, dense longitudinal neuroimaging (DLN) acquires multiple brain scans (typically MRI) of individuals’ timepoints across a short window of time to estimate individual trajectories of brain change (Booher et al., 2025).

Compared to neurotypical populations, little is known about the temporal-spatial brain dynamics of the language learning among individuals with communication disorders either developmentally or in the therapy setting due to the lack of brain imaging methodologies that are well suited to use with individuals with communication disorders (e.g., infants, young children, individuals with medical implants, etc.) in ecologically valid, face-to-face, naturalistic interactions. Although the high temporal resolution of EEG and MEG makes these technologies suitable for capturing millisecond-scale changes in spoken-language phonological, semantic, syntactic, and gestural cues (e.g., Hernandez et al., 2022), they have poor spatial resolution. Additionally, they are susceptible to mouth and hand movement artifacts, as well as facial expressions, gestures, and body language in natural interactions. Alternatively, functional MRI has a high field of view (FOV). However, the positive BOLD response in fMRI is very slow, beginning ∼500 msec and peaking ∼3-5 secs *after* the onset of a target event, making it poorly suited to capturing millisecond changes in the brain in response to the continuous millisecond fluctuations of the speech signal. The BOLD signal is also highly vulnerable to participant movement; thus, participants must remain as immobile as possible throughout the experimental task. Additionally, MRI scanner noise can be significant, making fMRI less well-suited for studies of spoken language. In contrast, the emerging field of optical functional near-infrared spectroscopy (fNIRS), and, in particular, recent advances in commercially available, compact, wearable high-density diffuse optical tomography (HD-DOT) brain imaging technology, allow participants to move naturally and interact spontaneously in naturalistic or clinical settings.

In this report, we use a high-density observation approach to examine the within-person learning trajectory of an individual participant over a series of eight sessions at two levels of description: (i) changes in performance, and (ii) changes in the hemodynamic response. For this report, we examined the participant’s ability to learn a set of spoken sentence stimuli using a sentence-repetition task. In children, sentence repetition tasks are not only part of the core subtests of the language assessment batteries (CELF, TOLD, etc.) but they have been proposed as a diagnostic marker for Developmental Language Disorder (DLD) in English, Catalan, and European Portuguese, and Cantonese-speaking children (Ahufinger et al., 2021; Archibald & Joanisee, 2099; Conti-Ramsden et al., 2001; Redmond et al., 2003; Stokes et al., 2006). In patients with head injury, the Sentence Repetition test (SRT; Spreen & Strauss, 1998) has been shown to be a reliable measure not only of the duration of unconsciousness but also of left-hemisphere injury (Meyers, Volkert, & Diep, 2000). Similarly, sentence repetition is a critical component of audiologic evaluations, such as speech audiometry (SRT, WRS), which evaluates the practical implications of hearing loss and influences diagnostic and treatment strategies (Billings et al., 2023; Naqvi & Sutton, 2025).

We also examined the pattern of the hemodynamic response in the frontal lobe. Sentence repetition performance has been viewed as a verbally mediated memory task and as a reflection of linguistic ability (e.g., MacDonald & Christiansen, 2002). It has also been shown to be a valid measure of processing effort among individuals with neurological disorders (Schroeder & Marshall, 2009). Among young adults with DLD, atypical levels of activation, spread of activation, and recruitment of frontal areas, as compared to controls, have been observed during verbal working memory tasks using fNIRS (Berglund-Barraza et al., 2019). Similarly, in functional MRI (fMRI) studies of DLD, atypical patterns of activation have been observed in frontal regions during both resting-state and verbal working memory tasks (Weismer et al., 2005).

For optical brain imaging to be a viable option for microgenetic studies, reliable within-person measures are needed that can differentiate the genuine hemodynamic response to the *learning trajectory* from generalized physiological fluctuations in oxygenated (HbO) and deoxygenated (HbR) hemoglobin levels. Although wearable fNIRS technology is comfortable, quieter than MRI, and can be used with children and those with implanted devices (i.e., cochlear implants), to be feasible in microgenetic studies, several additional criteria must be met. These include: 1) standardized probe array protocols to ensure consistent source-detector placement over multiple sessions, 2) standardized a priori signal optimization, average light levels across channels, and signal-to-noise ratio (SNR) levels to ensure the same signal quality levels are met over the sessions, 3) data processing algorithms than include superficial signal regression, which utilizes global variance of temporal derivatives (GVTD; Sherafati, 2020) index in the general linear models for censoring of high motion time points, 4) data processing algorithms that can provide researchers with both channel and brain space data, and 5) for the brain space analysis, algorithms that provide the same level of spatial resolution regardless of whether researchers use participants’ structural MRIs or an MNI database.

## 2. Methods

### 2.1 Data and code availability

Data and analysis scripts will be publicly available upon acceptance at our NeuroDOT repository, https://www.nitrc.org/projects/neurdot and on OpenfNIRS, https://openfnirs.org/.

### 2.2 Participant

The participant was a 63 year-old female who was strongly right-handed and a monolingual English speaker. The participant was not using psychotropic medications such as stimulants or anti-depressants and did not have a history of neurological disease, seizure disorder, psychiatric diagnosis, or comorbid conditions (e.g., Alzheimer’s, dementia, etc.). The participant had normal language, had never received a diagnosis of a language or learning disorder, and had never received language or learning therapy services. Structural MRI confirmed the absence of any neurological condition (e.g., stroke, tumor, etc.). The participant consented to the protocol in accordance with the Declaration of Helsinki and the guidelines of the University Institutional Review Board (IRB), which approved the protocols (IRB-24-258 and IRB-22-189).

### 2.3 Experimental design

The participant completed the same session a total of eight times over a period of nine weeks. To control for potential fatigue effects, each session started at ∼ 10 am each morning. During each session, the participant completed the same experimental paradigm consisting of two tasks: a spoken sentence repetition task and a spoken sentence comprehension task. The paradigm consisted of a total of 40 blocks (eight sentence repetition blocks, 12 sentence comprehension blocks, 20 auditory fixation blocks). Each block contained five sentences lasting ∼ 47 secs in duration. The auditory fixation blocks were 10 secs in duration for a total experimental paradigm time of 18 minutes. The blocks were presented in the same fixed-random order during each session. To ensure the participant understood the task, a series of practice sentence trials was conducted prior to the experimental task. The practice trials followed the same structure as the experimental trials but used different sentences from those presented during the experiment. The participant responded verbally to each sentence, and their responses were digitally recorded for offline accuracy assessment.

The experiment took place in a quiet, low-lit room. During each session, to reduce movement artifacts, the participant rested comfortably in a large King heavy-duty hydraulic “barber” chair with their legs on padded calf rests and their feet resting flat on the steel footrest. The chair was approximately four meters from two loudspeakers on the table in front of the participant at 0° azimuth. To further reduce potential upper-body movement, a large pillow lap desk was also placed on the participant’s lap, and they were asked to place their elbows on the lap desk and hold the microphone in both hands to stabilize their head and upper body. The participant’s responses were recorded using a Sennheiser e835 Live Vocal Microphone, positioned approximately 3 inches from their mouth, and captured on an Apple iPad. For the repetition task, the participant was asked to listen to a sentence, wait for the “beep,” and then repeat it. The beep consisted of a 500 Hz tone that occurred 1.5 secs after the end of each sentence.

### 2.4 Stimuli

The sentences in the repetition task were canonical Subject-Verb-Object, grammatically complex utterances comprised of one-syllable real words, where either only the first noun was animate (e.g., *the white owl bumped the broom under the very tall tree*”) or where the first two nouns were both animate (e.g., “*the black cat chased the bird into the very wet mud”*). Each sentence was digitally recorded individually at 44.1 Hz by a female English monolingual speaker. All the nouns and verbs ranged from two to four consonants in length. The verbs were all in the regular past tense form (e.g., *bumped, chased*). The nouns all had an age-of-acquisition (AoA) of ≤ 7 years 9 months (2 years 6 months – 7 years 9 months), and high concreteness (538 – 637), familiarity (453 – 624), and imageability ratings (499 – 638) (Kuperman, 2012; Walker & Hulme, 1999). To inhibit lexical cohort interference effects and because word frequency and phonotactic probability differentially influence the ability to hold words in memory (Coady et al., 2010; Mainela-Arnold, et al., 2010) all words were high frequency (≥ 100 MRC database). To prevent priming effects, a noun and a verb occurred only once in each block. The auditory fixation, “rest,” blocks consisted of audio recordings of nature sounds (e.g., birds) or Tibetan bells. To prevent habituation effects, each auditory fixation block was a different recording of the birds or Tibetan bells. The stimuli were presented to the participant at a comfortable listening level, which remained constant across the eight sessions.

### 2.5 fNIRS system and data acquisition

To measure brain activity, a NIRSport2 (NIRx, Medical Technologies LLC, Berlin, Germany), portable, wearable, continuous-wave, multi-channel NIRS system was used to acquire hemodynamic changes in the prefrontal cortex while the participant completed the tasks. The prefrontal probe array consisted of eight emitter and seven detector optodes at source-detector distance of three cm, providing 20 channels in total (Figure 1A). The eight LED emitters consisted of dual-tip NIRS optode light sources, emitting near-infrared light at 760 and 850 nm, with the sources having a maximum light levels of 25 mW_max_ per wavelength.

**Figure 1.**
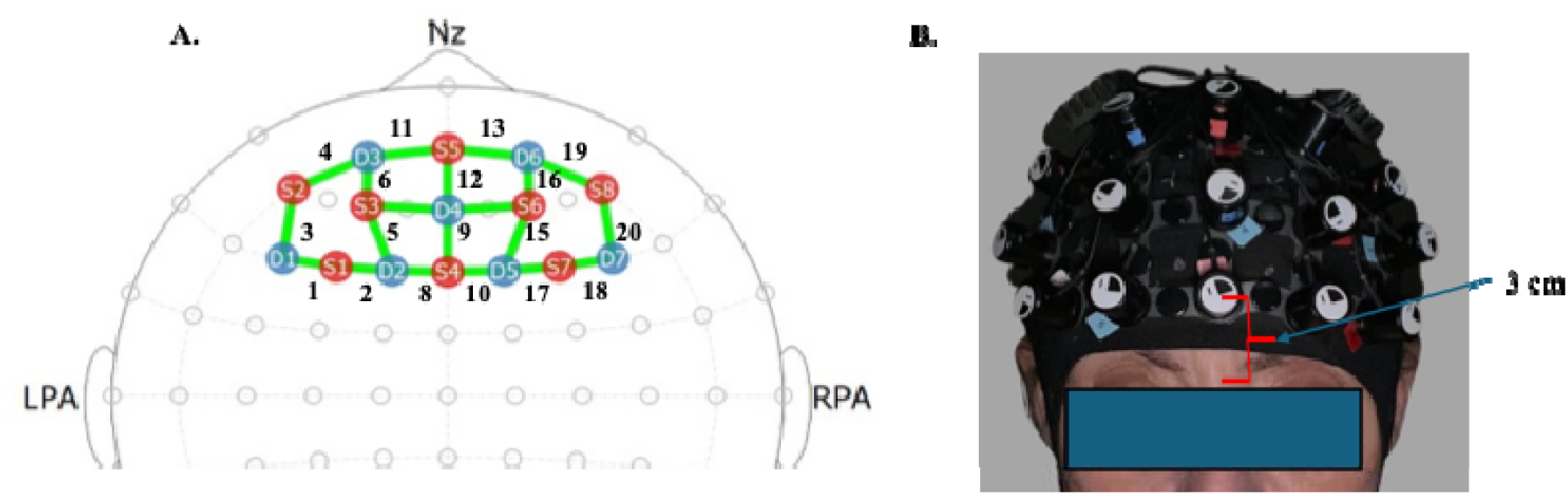
**A.** NIRSport2 7×8 Optode layout. Emitters are red and detectors are blue. Green lines represent channels. **B.** Example of cap placement and 3 cm distance from nasion to Source 5(S5) Fpz.

The optodes were placed bilaterally and symmetrically on the participant’s forehead using an NIRScap spandex fNIRS cap (NIRx, Medical Technologies LLC, Berlin, Germany) marked according to the 10-20 system to guide and standardize optode placement for each session. Before the cap fit, the participant’s hair was combed away from the forehead and secured to prevent it from obscuring the optodes. The participant’s head circumference, distance between right to left preaural area, and nasion to inion were measured. An NIRScap fitted with the array of sources and detectors was then placed on the participant’s head. To ensure consistent array placement across sessions, cap placement was standardized by ensuring that S5 was 3cm directly above the nasion at Fpz (See Figure 1B.). Signal calibration and recording of fNIRS data was completed using Aurora 1.4. Acquisition Software (v2021.9, Medical Technologies, Berlin, Germany). If the signal quality, as indicated by Aurora Software, met the a priori criteria, the task was administered. Data were recorded at a sampling frame rate of 10.17 Hz.

### 2.6. Data processing, image reconstruction, and analysis

Data were processed using the NeuroDot (https://www.nitrc.org/projects/neurdot) processing pipeline in MATLAB (Eggebrecht & Culver, 2019; Speh, et al., 2025). Data analysis and image reconstruction occurred in five steps: light-level measurement pre-processing, anatomical light modeling, image reconstruction, spectroscopy, and spatial normalization. For details regarding these procedures see Sherafati et al., (2020) and Eggebrecht et al., (2014).

#### 2.6.1 Measurement pre-processing

The raw .snirf files were converted to .mat files and pre-processed using a standardized NeuroDOT pre-processing pipeline. Prior to starting the data pre-processing steps, NeuroDOT general data quality assessment visualizations were used to assess cap light levels and raw data quality for all eight sessions (See Figure 2). Sessions were included in final analysis if they met the following criteria: 1) good coupling across the entire cap as evidenced by light levels being close to the maximum NIRSport2 of 25 mW_max_ per wavelength (e.g., white and yellow diamonds below) across channel measurements (Figure 2A), 2) high signal-to-noise ratio (SNR) for cardiac pulse band of frequencies (Figure 2B), 3) no optodes having bad measurements (Figure 2C), and 4) all source and detector meeting a threshold of 6% based on the histogram of signal percent standard deviation (Figure 2D).

**Figure 2.**
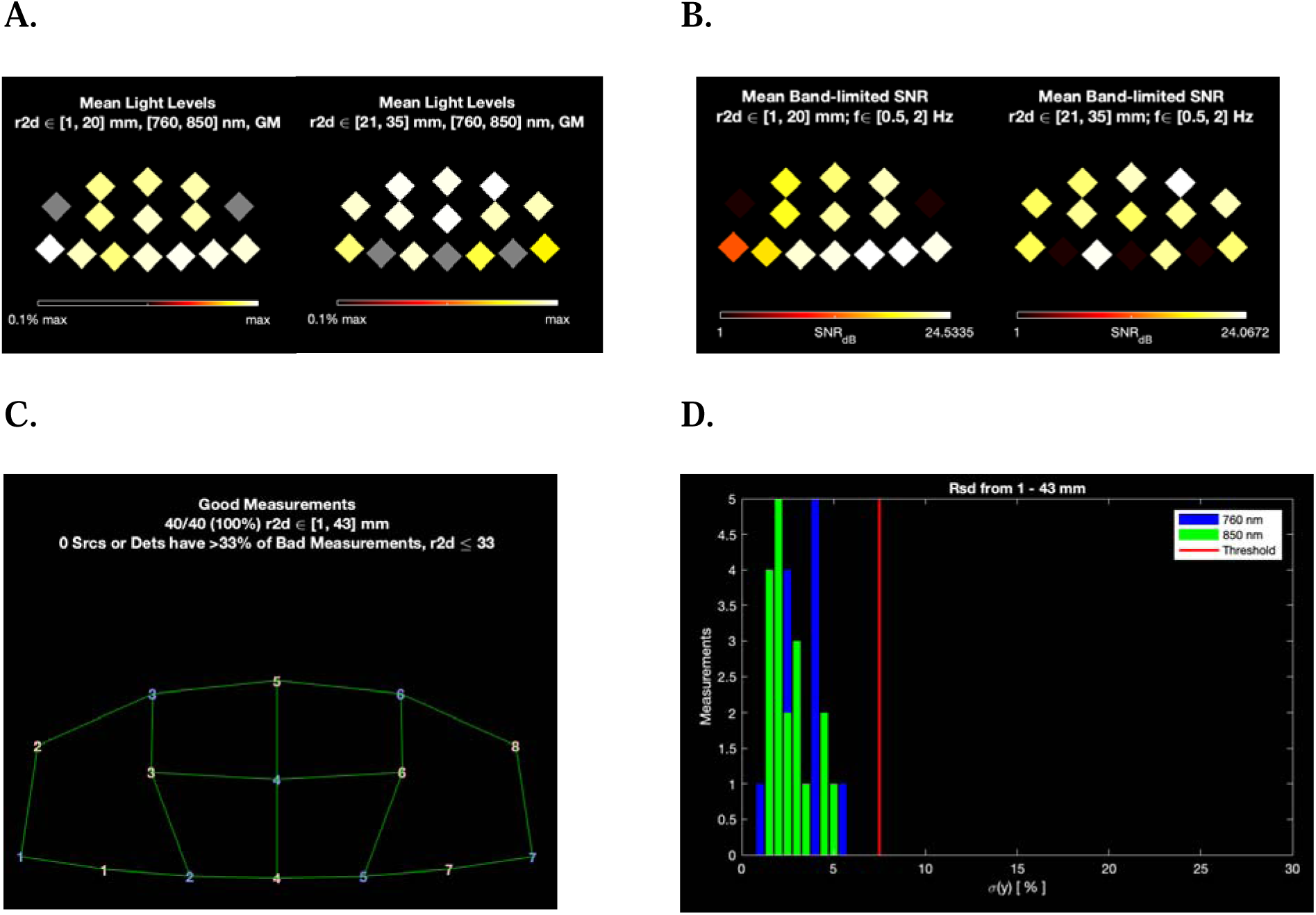
NeuroDot raw data quality assessment. **A.** Average light levels across channel measurements. **B.** signal-to-noise ratio (SNR) for cardiac pulse frequency band. **C.** Optode quality measurements. **D.** Histogram of signal percent standard deviation all source and detector met a threshold of 6%.

All sessions that met the inclusion criteria were then pre-processed. A visualization of select preprocessing steps can be found in Figure 3. Figure 3A visualizes the average wavelength spectrum, including the cardiac pulse and stimulus frequencies. Before filtering the data, time-series logmean light levels were derived from baseline mean raw source-detector (channel) data across the entire experiment duration. Any measurements with a temporal standard deviation greater than 7.5% were excluded to avoid data contamination from non-physiological variance cause by motion artifact. Data was then fast Fourier transformed to show the presence of periodic, or nearly periodic waveforms in the measurements. This clearly showed the cardiac pulse with a period of approximately one second and appears as a spike at a frequency of approximately 1Hz (Figure 3B). The remaining measurements are detrended and high-pass filtered at 0.01 Hz to remove any linear trend or drift in the measurement. Figure 3C shows the effect of this initial filter as frequencies below the cutoff frequency are significantly attenuated (frequencies higher than the cutoff pass through the filter). Note that the cardiac pulse and stimulus frequencies are still present in the spectrum. The spectrum (Figure 3D) shows attenuation of frequencies above the cutoff frequency (frequencies lower than the cutoff pass through the filter). Next, a common signal was estimated from the superficial measurements and regressed out of all measurements. Figure 4E shows a much stronger peak at the stimulus frequency relative to other frequencies. An additional low pass filter attenuates frequencies above 0.1 Hz to ensure the entirety of the heartrate is excluded from the stimuli. The second low pass filter drastically reduces the high-frequency noise, leaving only waveforms at the stimulus frequency (Figure 3F). Data are resampled at 1 Hz (Figure 3G). Finally, measurements are conservatively block averaged based on block duration, 47 seconds for repetition blocks and 9 seconds for auditory fixation blocks.

**Figure 3.**
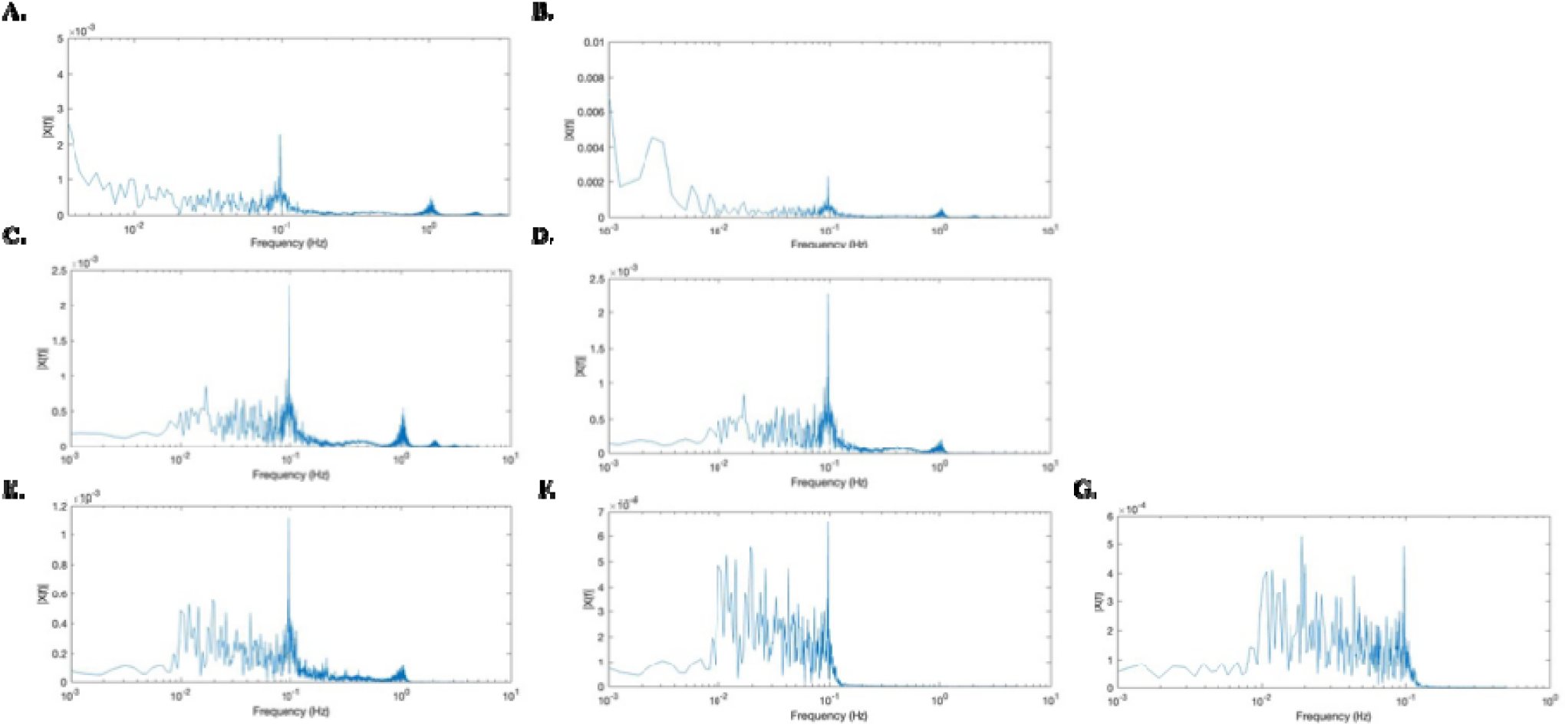
NeuroDot preprocessing pipeline. **A.** No filtering **B.** Log mean and detect noisy channels **C.** Detrend and highpass filtered at .01 Hz **D.** Low pass filtered at 1 Hz. **E.** Superficial signal regression. **F.** Second low pass filter at .1 Hz. **G.** Resampling at 1 Hz.

**Figure 4.**
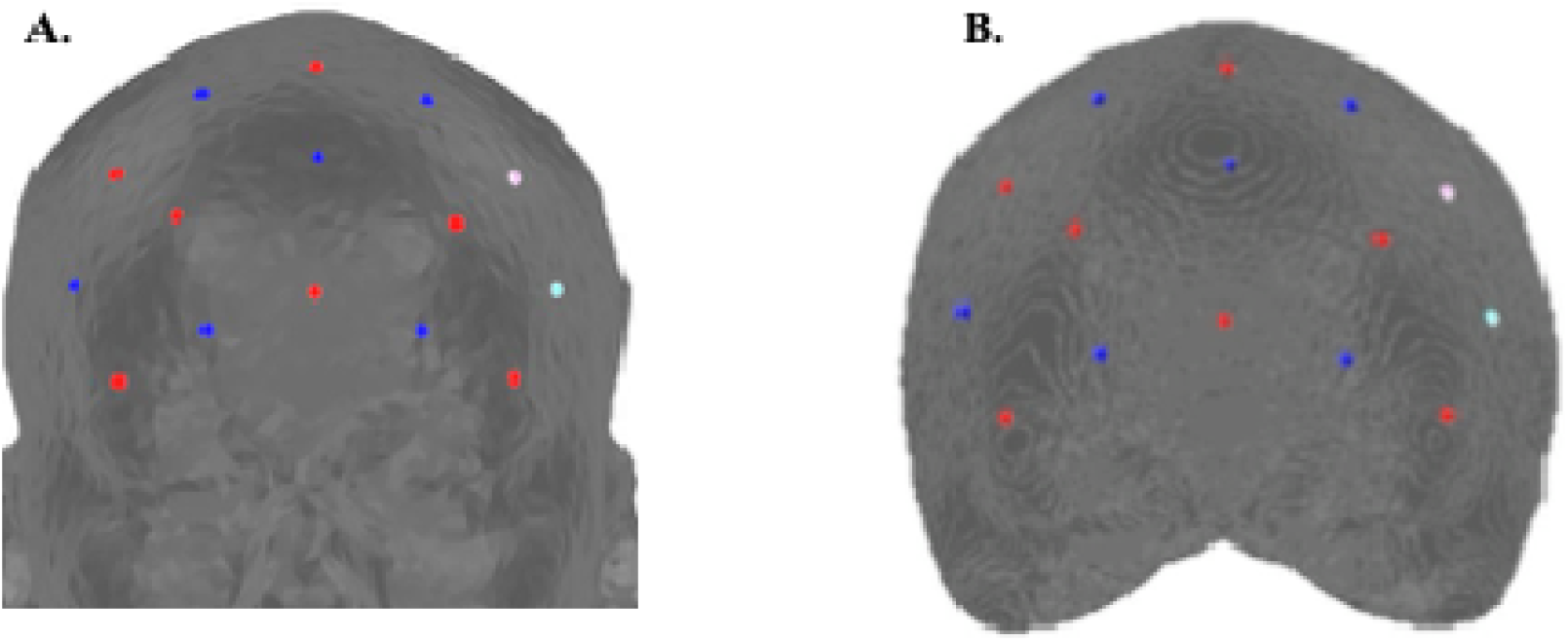
**A.** Subject MRI low density mesh **B.** Subject MRI high density mesh

The spectral plots were used to determine the participant’s average heart rate for each session and to ensure that the low pass filter values that were used excluded the cardiac pulse. The presence of periodic, or nearly periodic waveforms in the measurements show the cardiac pulse with a period of approximately one second and appears as a spike at a frequency of approximately 1Hz (Figure 3A-E) with the harmonics being clearly visible due to the filter in Figure 3C.

#### 2.6.2 Channel-space data processing

Following the pre-processing pipeline, changes in oxygenated hemoglobin (ΔHbO_2_) and deoxygenated hemoglobin (ΔHb) concentrations were derived based on the Modified Beer-Lambert Law with a partial path-length factor of 1.0 for both wavelengths. Global variance of temporal derivatives (GVTD) index was used to detect and censor motion artifacts in the data (Sherafati et al., 2020). A general linear model was then conducted on the channel motion-free data. The block-averaged ΔHbO_2_ and ΔHb concentrations for each of the 20 channels for the Sentence Repetition and Auditory Fixation conditions were used in the analysis.

#### 2.6.3 Brain- space: MRI segmentation

An MRI scan was also performed at the Center for BrainHealth, a research center under the University of Texas at Dallas. A 3T Siemens Magnetom Prisma scanner with a 32-channel head coil was used. High-resolution anatomical images were acquired using a transverse MPRAGE T1-weighted sequence with the following parameters: TR = 2300ms; TE = 2.26ms; flip angle = 8°; acquisition matrix = 256 x 256; voxel size = 1 mm^3^, slices = 208, duration = 5:21 mins. Freesurfer was used to create a model of the participant’s brain from the MRI data.

#### 2.6.4 Brain- space: Anatomical light modeling, image reconstruction, spectroscopy, spatial normalization, and data visualization

The first step in producing a head model, is ensuring that the NIRSport2 prefrontal probe array spatial and topological information was transferred to a pad file, which is NeuroDOT’s representation of the imaging system’s optode array positions. NeuroDOT’s pad file generation pipeline was utilized. Anatomical light modeling requires either a subject-specific or an atlas-based segmented volume. In this study, we reconstructed the image using the participant’s structural MRI. Using the pad file as an input, a mesh (3D representation) was generated and used to efficiently align the optical array to its position on the 3D head model using Nirfaster (Dehghani et al., 2008. NeuroDOT then leverages AlignMe, its app for array placement, to efficiently scale and relax the mesh onto the head model to reflect accurate participant probe placement (Figure 4A). Methods for the spring-relaxation energy minimization algorithm used are described in Joseph et al., 2006. Next, a smaller high-density mesh is generated based on optode location to reflect the coverage of the probe array (Figure 4B).

Lastly, using NIRFASTer (Dehghani et al., 2008), a sensitivity matrix was generated for each source detector measurement (see Eggebrecht et al., 2014; Ferradal et al., 2014 for a full description of methods used). Absorption image volumes were reconstructed from the processed measurements based on a regularized inversion of the sensitivity matrix (for details, see, Eggebrecht et al., 2012). The sensitivity matrix was inverted using Tikhonov regularization (λ = 0.01, λ = 0.1, gSigma = 3), and spectroscopy was performed using literature-derived values (Eggebrecht et al., 2012; Bluestone et al., 2001). These steps produce volumetric time-series data of relative changes in oxyhemoglobin (ΔHbO_2_), deoxyhemoglobin (ΔHbR), and total hemoglobin (ΔHbT) concentrations (Bluestone et al., 2001). For the brain space data, following the pre-processing and reconstruction of the data, a general linear model was also conducted on the motion-free data. The data visualization pipeline was then completed to spatially normalized the participant’s MRI to MNI space to be visualized on a surface model of the cortex. To compare across days, brain maps were displayed using a fixed color scale applied uniformly. A scale value of 5e-05 was selected to provide stable visualization and was not adjusted on a per-map basis.

### 2.7 Statistical Analysis

For behavioral performance, a dense word-level analysis of the participant’s repetition performance was conducted, and a repeated-measures ANOVA examined word-level accuracy across sessions (Redmond, 2003). For channel-based analysis of the hemodynamic response, a repeated-measures ANOVA was first conducted to examine channel-wise hemodynamic change across the prefrontal cortex for the Auditory Fixation condition, to determine whether the hemodynamic response was stable across the five sessions. Once it was determined that the channel-wise mean changes in oxygenated hemoglobin (ΔHbO_2_) and deoxygenated hemoglobin (ΔHb) concentrations did not differ for the Auditory Fixation condition across the sessions, channel-wise mean changes in ΔHbO_2_ and ΔHb concentrations were then examined for the Repetition condition using a GLM analysis with Auditory Fixation and channel as covariates. Data were corrected using the Greenhouse-Geisser correction applied to the probability values to adjust for repeated measures.

## 3. Results

### 3.1 Signal quality and heart rate measures

Only sessions one through five met all inclusion requirements. Although sessions six and seven had good cap coupling, high signal-to-noise ratio, and good optode measurement, they did not meet our 6% signal quality thresholds. Session eight met all the inclusion requirements, but because the poor signal quality of sessions six and seven created a gap in our ability to track the microgenetic changes in the brain and behavior patterns of the participant’s learning trajectory for this report, only sessions one through five were included in the subsequent analysis (See the Supplemental Material 1 for the NeuroDOT raw data quality assessment figures). The participant’s average heart rate for each session was identified from the NeuroDOT pre-processing pipeline and low and high pass filters were based on these values (see Supplemental Material 2 for the spectral plots for the first five sessions).

### 3.2 Behavioral Performance

As shown in Table 1, the participant’s accuracy over the course of the first five sessions was high (96.43% - 97.44%) and did not differ over the sessions *F*(4, 35) = .285, p = .88, partial eta ^2^ = .032, power = .10. However, although accuracy was high and did not change over the sessions, one-way *t* tests (one-tailed) revealed that, with the exception of session 2, the participant’s ability to accurately repeat the sentences was significantly different from 100% (See Table 1). Thus, despite repeating the same sentences in the same order over the course of the 5 sessions, the participant’s performance did not improve, indicating that the participant did not remember nor learn the stimuli.

**Table 1.** Accuracy, *t*-value, *p* -value and Cohen’s *d* for the precent words repeated accurately for the first 5 sessions.

| Session | Accuracy | SD | $t$ | $p$-value | Cohen's $d$ |
| --- | --- | --- | --- | --- | --- |
| 1 | 96.42 | 2.28 | -4.434 | 0.002 | 2.28* |
| 2 | 95.63 | 7.44 | -1.663 | 0.07 | 7.44 |
| 3 | 95.11 | 2.87 | -4.812 | 0.001 | 2.87* |
| 4 | 96.42 | 5.35 | -1.998 | 0.043 | 5.34* |
| 5 | 97.44 | 3.62 | -2.285 | 0.028 | 3.61* |

\*Significantly different from 100%

Similarly, the participant’s performance also did not improve over the course of the eight blocks within each session. This lack of improve indicates that there was no learning effect either within or across the sessions. As seen in Figure 5, changes in the participant’s performance over the blocks also shows that no block of sentences were inherently easier for the participant to remember. Instead, the participant’s performance fluctuated over the course of the eight blocks in a different pattern from Day 1 to Day 5.

**Figure 5.**
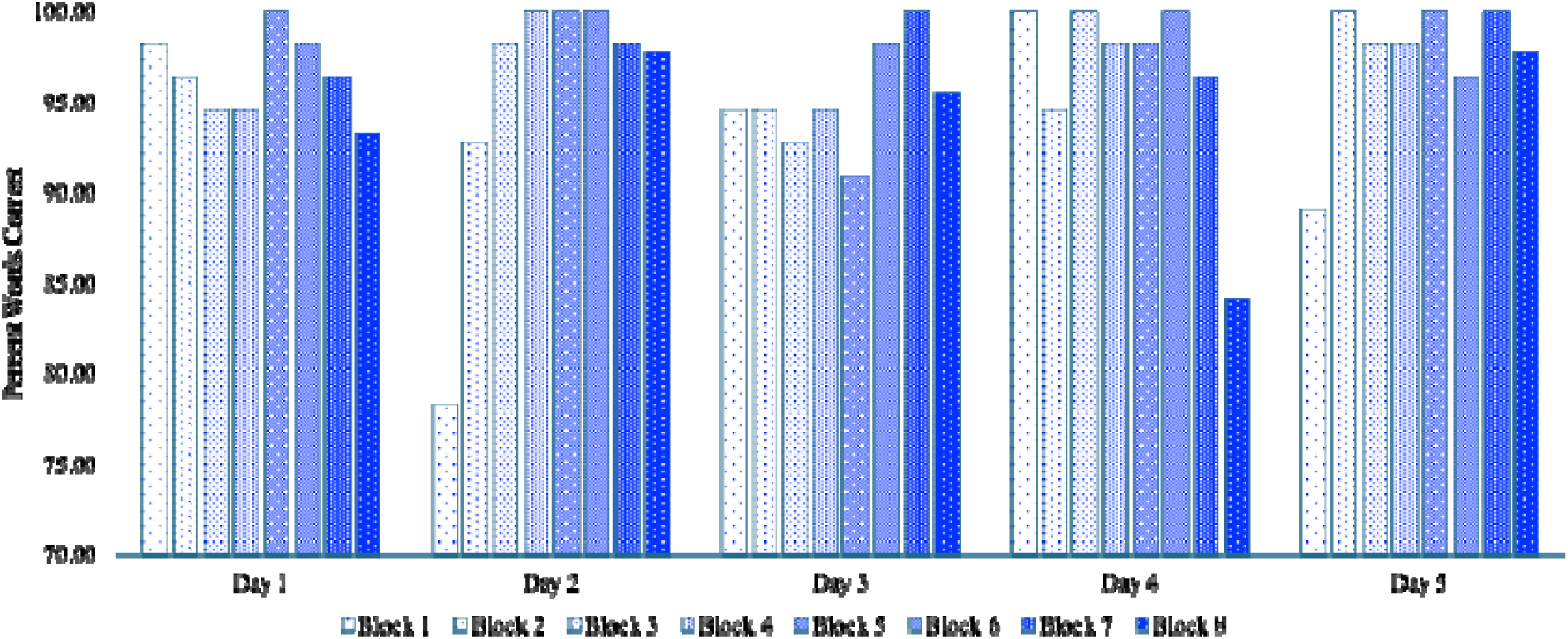
Percent Words Correct for each block of the Repetition task for Day 1 through Day 5.

### 3.3 Channel-based analysis of Hemodynamic response

#### Auditory Fixation

A repeated measures 2 (ΔHbO_2,_ ΔHb) x 5 (session) general linear model with channel as a covariate revealed that although the average concentration for the Auditory Fixation condition did not differ statistically over the five sessions *F*(2.3, 85.2) = .76, *p* = .48 partial eta ^2^ .02, power .24, there was a main effect of ΔHbO_2_ and ΔHb concentration *F*(1, 37) = 10.59, *p* = .002 partial eta ^2^ = .22, power = .88, and a session by ΔHbO_2_ and ΔHb interaction effect *F*(2.3, 85.2) = 5.73, *p* = .003 partial eta ^2^ .134, power .88. Post hoc analysis revealed that ΔHbO_2_ did not differ statistically for the Auditory Fixation across the first five sessions *F*(1.92, 37.78) = 2.68, *p* = .09 partial eta ^2^ = .136, power = .486 nor by hemisphere *F*(2, 17) = .55 *p* = .586, partial eta ^2^ = .061, power = .126, nor was there a session by hemisphere interaction *F*(3.85, 32.78) = 1.48, *p* = .23, partial eta ^2^ = .14, power = .40. In contrast, post hoc analysis revealed that ΔHb concentration did differ significantly for the Auditory Fixation over the five sessions *F*(2.86, 48.68) = 7.05, *p* < .001 partial eta ^2^ = .29, power = .96, but did not differ by hemisphere *F*(2, 17) = .24 *p* = .78, partial eta ^2^ = .028, power = .082, nor was there a session by hemisphere interaction *F*(5.728, 48.68) = 2.15, *p* = .07, partial eta ^2^ = .123 power = .69. Because ΔHbO_2_ concentration did not differ for Auditory Fixation for the first five sessions, the rest of the analyses were conducted for ΔHbO_2_ only.

#### Repetition

A GLM model with auditory fixation as a covariate revealed a main effect for session *F*(4,84) = 3.36, *p* = .013, partial eta ^2^ = .13, power = .82 where Δ*HbO_2_*concentration for the Repetition task differed significantly over the five sessions as well as by hemisphere effect *F*(2, 84) = 3.809, *p* = .026, partial eta ^2^ = .083, power = .67 where Δ*HbO_2_* concentration differed in the left, right, and center channels (See Figure 6).

**Figure 6.**
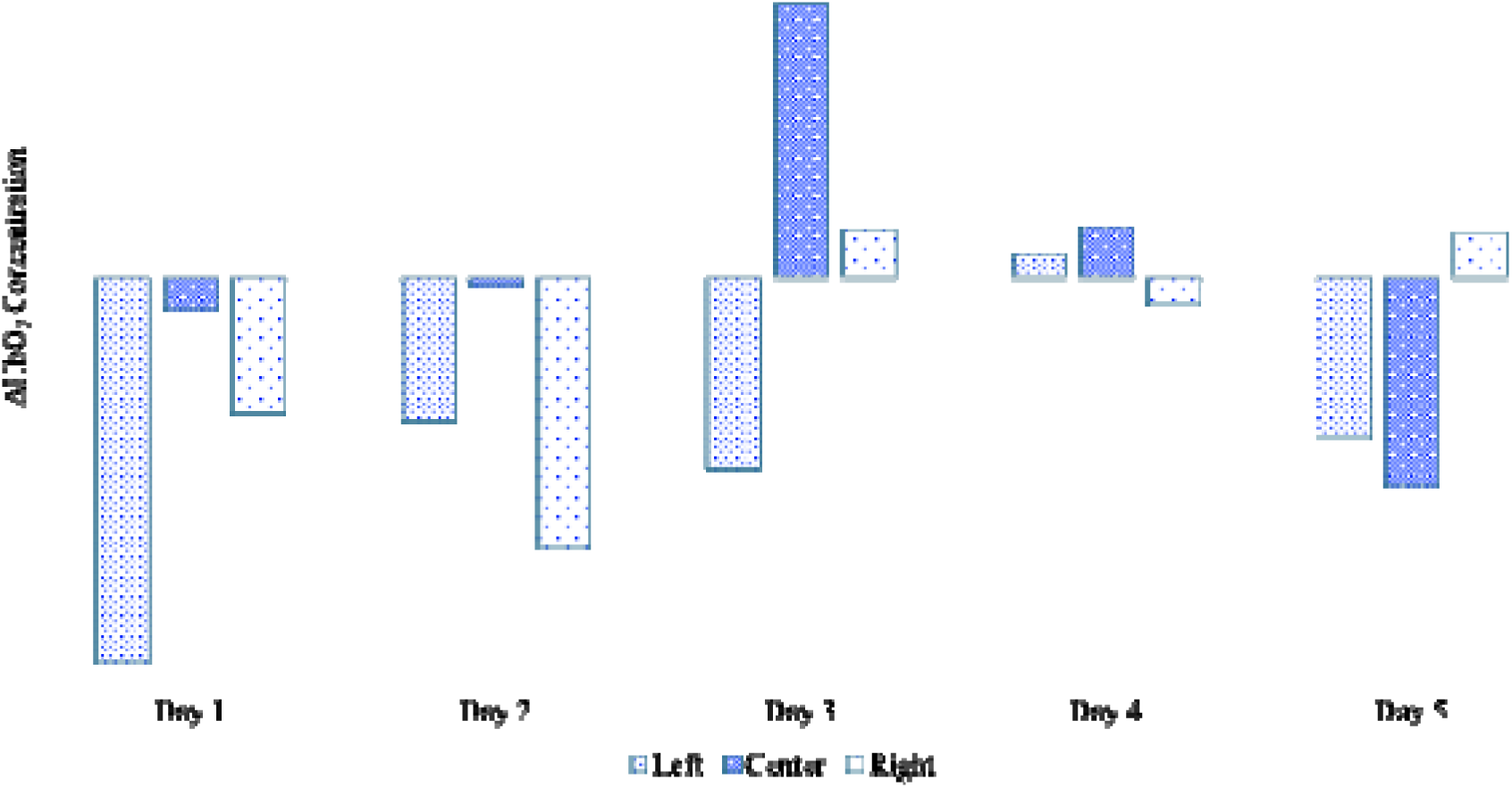
ΔHbO_2_ concentration for the Left, Center, and Right channels for the Repetition task.

### 3.4 Brain- Space visualization of Hemodynamic response

The ΔHbO_2_ concentrations in the prefrontal region for the average hemodynamic response for the auditory fixation blocks (baseline) subtracted from the average hemodynamic response for the repetition blocks for each of the five sessions can be seen mapped onto the participant’s structural MRI spatially normalized to MNI space in Figure 7.

**Figure 7.**
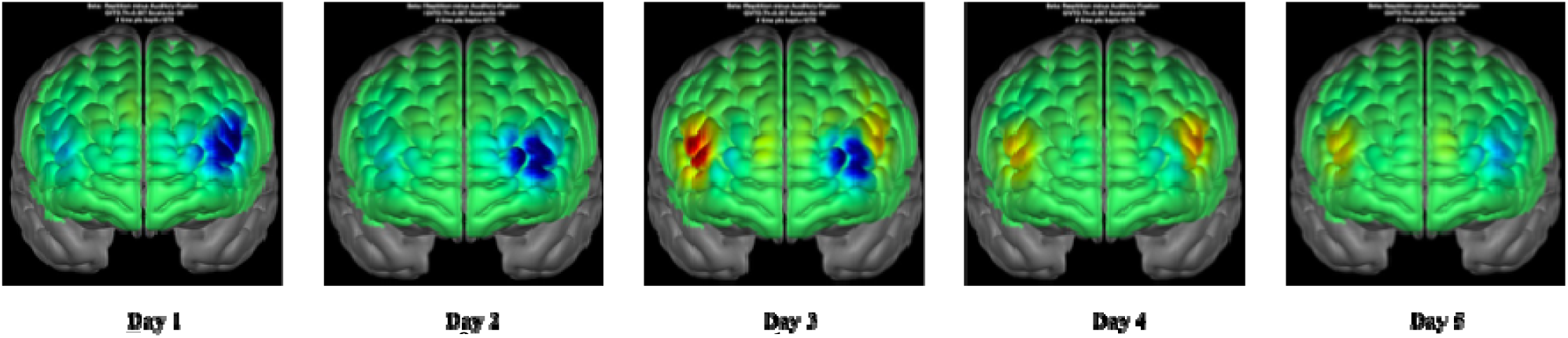
ΔHbO_2_ visualized for Repetition > Auditory Fixation for Day 1 through Day 5.

The figure displays the area of activations during the Repetition>Auditory Fixation which aligns with the channel-based statistical data showing that Δ*HbO_2_* concentration for the Repetition task differed by hemisphere over the five sessions, but shows more clearly that despite completing the same task over the course of the 5 days, the pattern of frontal activation was similar for Day 1 & 2, but *differed* qualitatively for Days 3-5.

## 4 Discussion

These findings demonstrate how wearable functional near-infrared spectroscopy (fNIRS) optical imaging methods, when combined with optical brain imaging data processing algorithms and dense-sampling research designs, can provide unique insights into the changing dynamics of brain and behavior over the course of a microgenetic study. We anticipated “learning,” as evidenced by an increase in sentence repetition performance over the course of the sessions as the participant began to remember the sentence stimuli. Instead, the participant’s performance was high but consistently statistically significantly different from 100% across sessions suggesting that in this case, the task was a stable measure of verbal working memory, regardless of how familiar the participant was with the task stimuli. We also expected a stable hemodynamic response as measured by the Channel-based analysis of ΔHbO_2_ concentration and the Brain-Space visualization of the Hemodynamic response. Although we did observe that the ΔHbO_2_ concentration remained stable for the Auditory Fixation condition and did not change over the course of the sessions, for the Repetition condition the pattern of ΔHbO_2_ concentration changed over the sessions. The visualization of the brain-space data revealed, remarkably, that the pattern of frontal region engagement by the participant while completing the *same* sentence repetition task over the five days changed significantly, with no two days being the same. Because the hemodynamic response for Auditory Fixation did not change over the five days, this suggests the presence of meaningful within-person differences in frontal engagement during the repetition task across the sessions.

The Brain-Space visualization afforded by NeuroDOT (https://www.nitrc.org/projects/neurdot) added unique information not available solely from the Channel-space analysis. Notably, NeuroDOT allowed for high spatial resolution brain maps that we were able to register onto the participant’s MRI. This suggests that, even though NeuroDOT was created with high-definition optical systems, when combined with the new high-definition wearable optical imaging technologies, new processing algorithms such as NeuroDOT can provide valuable spatial resolution on the order of fMRI. Therefore, even though the data for this report were collected using a sparse NIRSport 2 probe array without short channels, NeuroDOT was able to provide valuable spatial resolution.

This technical report also shows that behavioral assessments do not give us a complete picture about what is happening in speech repetition tasks. This has implications for audiology, as the standard diagnostic protocol consists of word and sentence repetition tasks. In speech and hearing, we focus on what is being heard and produced, rather than on what the brain is doing with this information. The imaging data from this study showed that the participant was engaging the frontal networks differently each session. This indicates that optical imaging techniques have the temporal and spatial resolution to track microgenetic changes in the learning strategies employed by speech, language, and hearing patients during diagnostic assessment and over the course of therapy. By leveraging neuroimaging afforded by wearable optical imaging systems, clinicians and researchers can now monitor changes in brain regions engaged within and across clients to identify individual differences in brain activation patterns, as well as patterns that may uniquely identify and discriminate disordered populations from neurotypical controls.

## Acknowledgments

This research was conducted using the data collected by the investigators. We are grateful for their and the participant’s contributions. We are also grateful for the support provided by the National Institute on Deafness and Other Communication Disorders K18 DC021149 (PI Evans) and the University of Texas Dallas Friends of Brain Health Grant (PI Skolasinska). The analyses and interpretations presented in this study are solely those of the authors.

