## Supplement 1 for "Highlighting the strengths of new wearable fNIRS, precision accuracy microgenetic research designs, and new NeuroDOT processing algorithm to measure change over time in clinical populations for Speech-Language and Hearing Researchers"

Supplemental 1

Data Quality assessments for each Session

A. Average light levels across channel measurements and signal-to-noise ratio (SNR) for cardiac pulse frequency band.

**Session 1:** 05_09: Passed

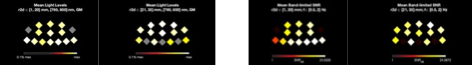

**Session 2:** 05_27: Passed

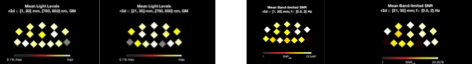

**Session 3:** 06_24: Passed
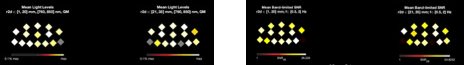

**Session 4:** 06_25: Passed

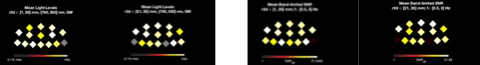

**Session 5:** 07_01: Passed

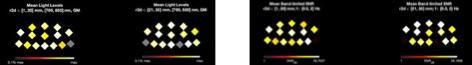

**Session 6:** 07_02: Passed
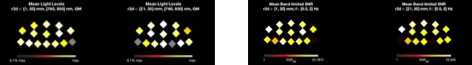

**Session 7:** 07_08: Passed

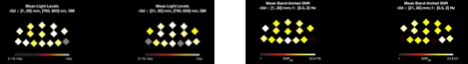

**Session 8**: 07_09: Passed

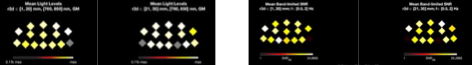

**B.** Optode quality measurements and histogram of signal percent standard deviation all source and detectors

**Session 1:** 05_09: Passed

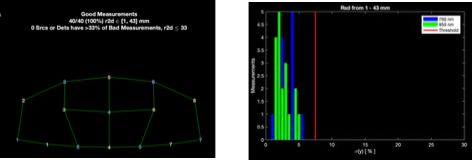

**Session 2:** 05_27: Passed

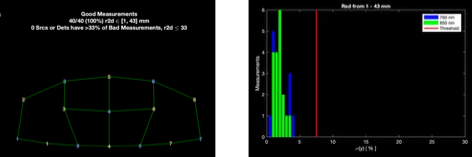

**Session 3:** 06_24: Passed

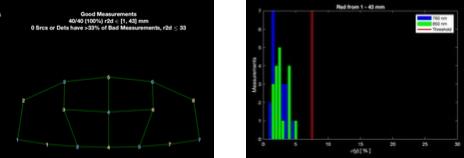

**Session 4:** 06_25: Passed

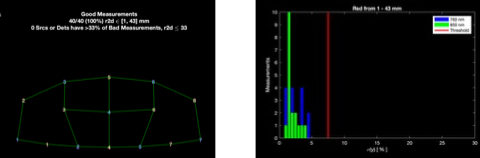

**Session 5:** 07_01: Passed

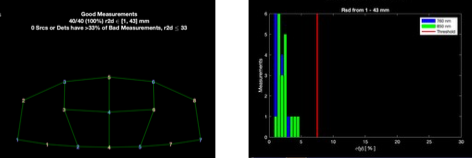

**Session 6:** 07_02: Did not pass signal threshold of 6%

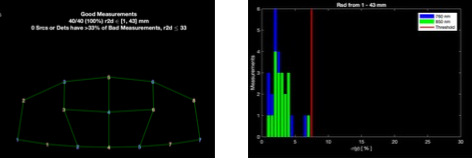

**Session 7:** 07_08: Did not pass signal threshold of 6%

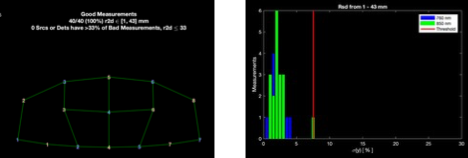

**Session 8**: 07_09: Passed

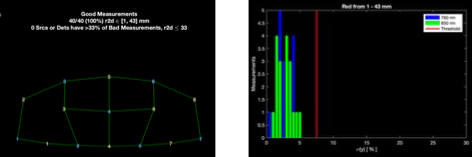
