## Supplement 2 for "Highlighting the strengths of new wearable fNIRS, precision accuracy microgenetic research designs, and new NeuroDOT processing algorithm to measure change over time in clinical populations for Speech-Language and Hearing Researchers"

Supplemental 2

**A.** Raw signal prior to processing for sessions one through five showing the average wavelength spectrum, including the cardiac pulse and stimulus frequencies

**Session 1:** 05_09

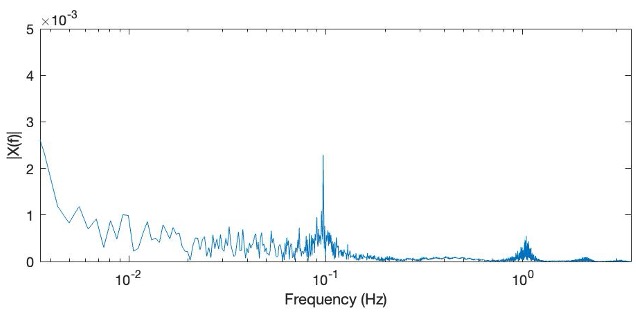

**Session 2:** 05_27

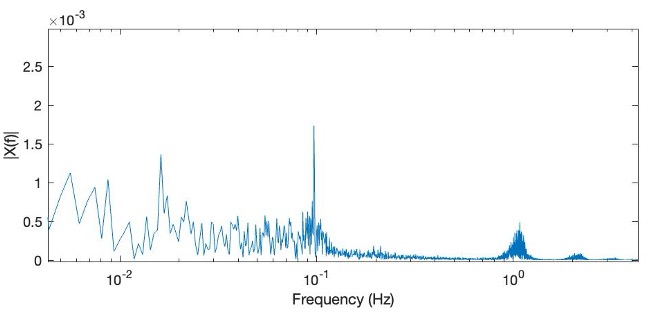

**Session 3:** 06_24

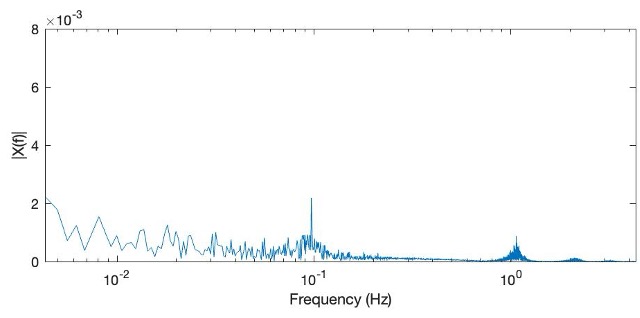

**Session 4:** 06_25

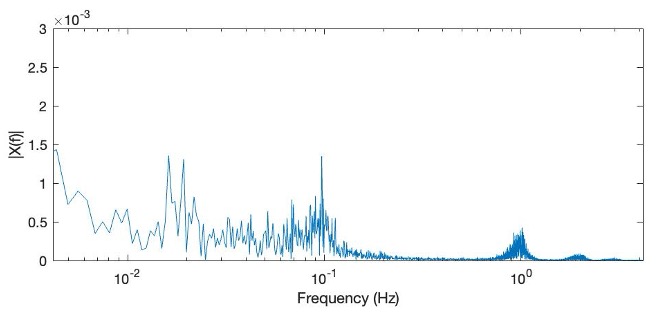

**Session 5:** 07_01

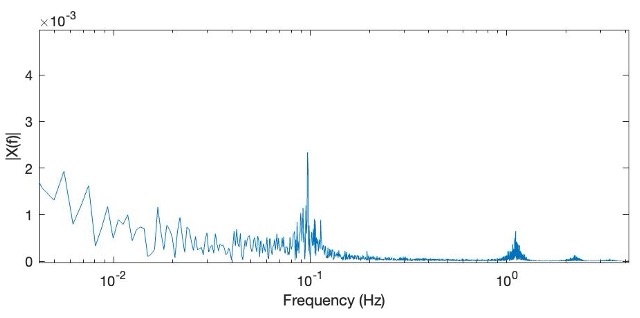

**B.**  Any measurements in the data from sessions one through five with extensive noise were excluded. Measurements were fast Fourier transformed to show nearly period waveforms that display the cardiac pulse with a period of approximately one second and appears as a spike at a frequency of approximately 1Hz.

**Session 1:** 05_09

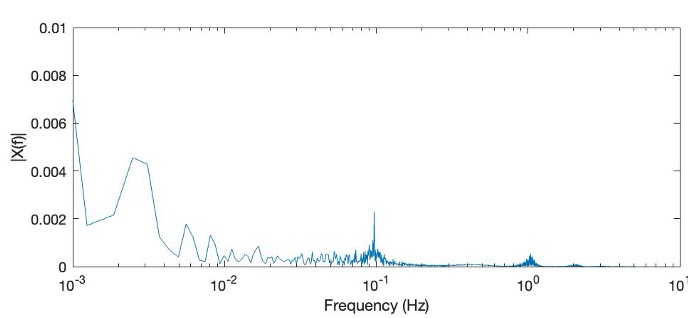

**Session 2:** 05_27

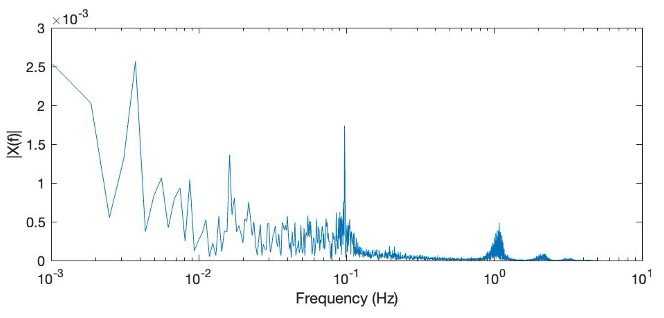

**Session 3:** 06_24

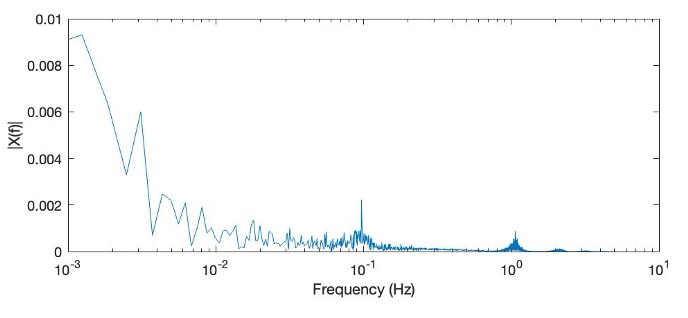

**Session 4:** 06_25

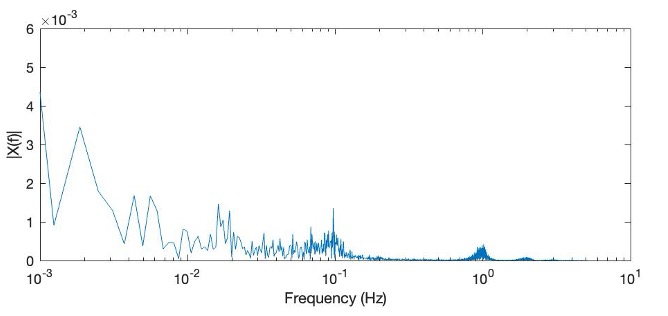

**Session 5:** 07_01

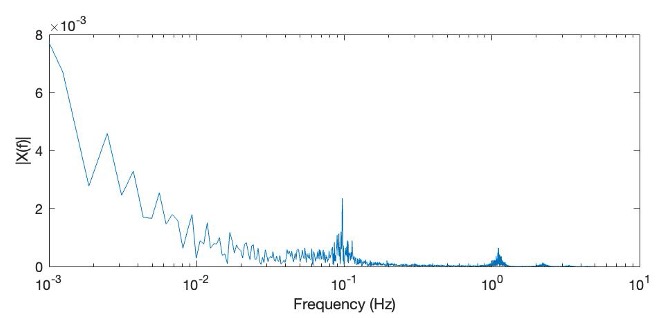

**C.** The remaining measurements in session one through five are detrended and high- pass filtered at 0.01 Hz to remove any linear trend or drift in the measurement. Note that the cardiac pulse and stimulus frequencies are still present in the spectrum.

**Session 1:** 05_09

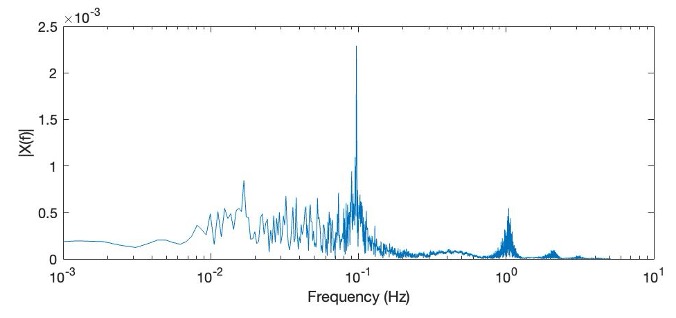

**Session 2:** 05_27

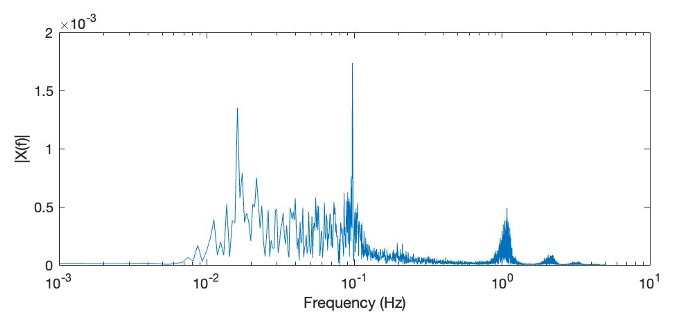

**Session 3:** 06_24

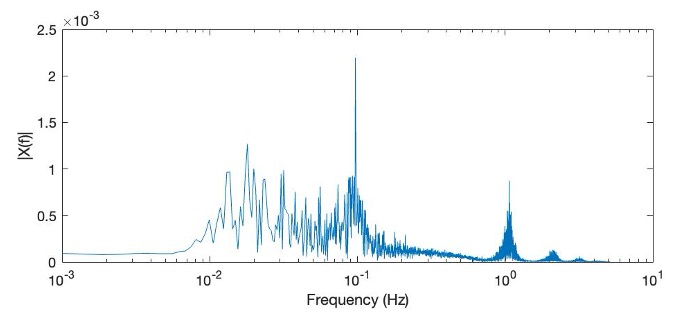

**Session 4:** 06_25

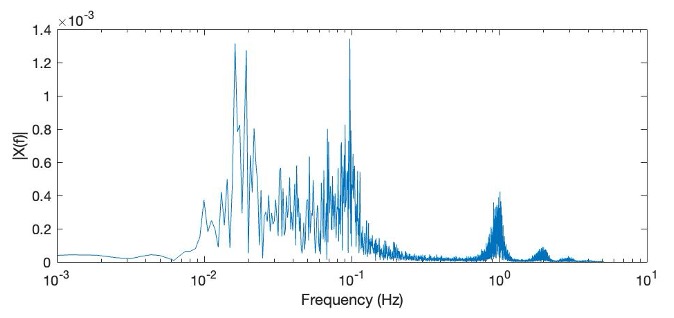

**Session 5:** 07_01

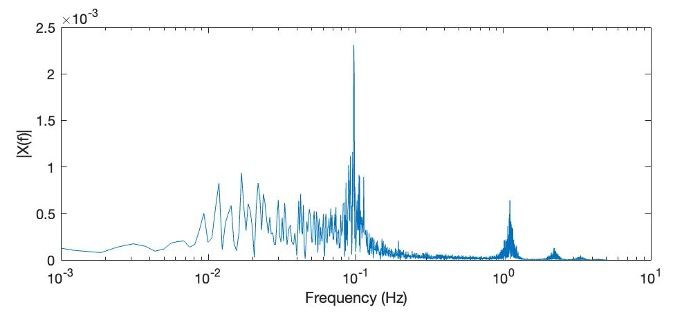

**D.** Remaining measurements from sessions one through five are low- pass filtered at 1 Hz.

**Session 1:** 05_09

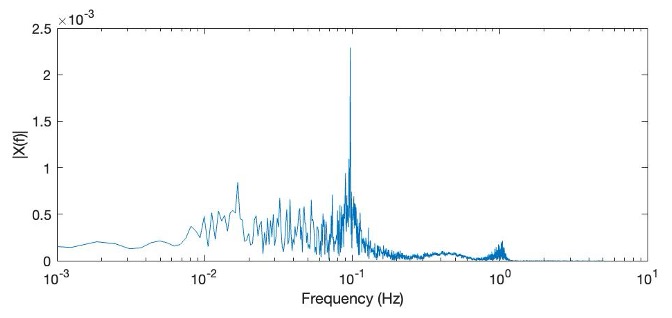

**Session 2:** 05_27

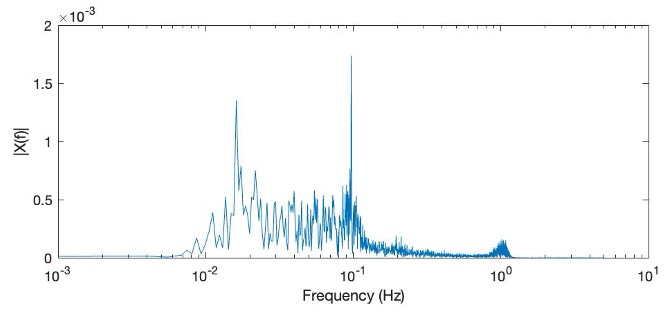

**Session 3:** 06_24

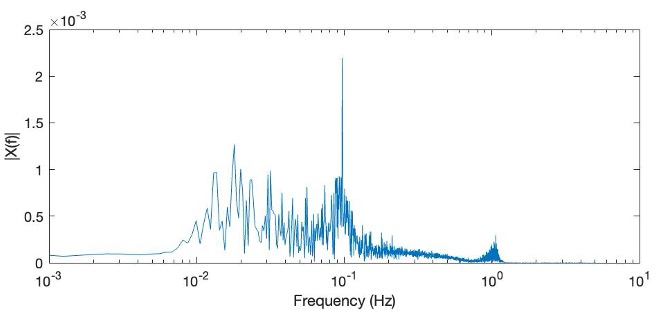

**Session 4:** 06_25

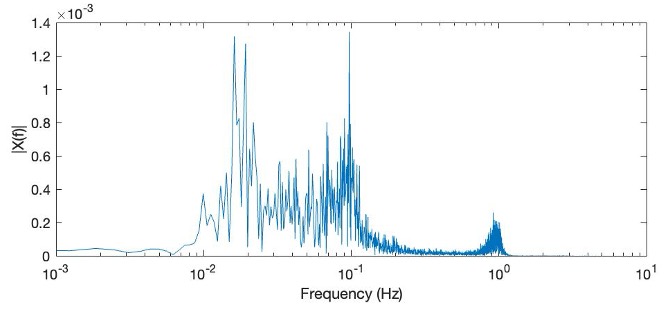

**Session 5:** 07_01

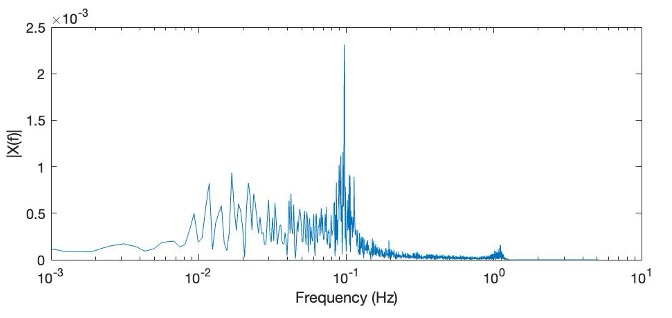

**E.** Remaining measurements from sessions one through five undergo superficial signal regression. Now the stronger peak at the stimulus frequency relative to other frequencies.

**Session 1:** 05_09

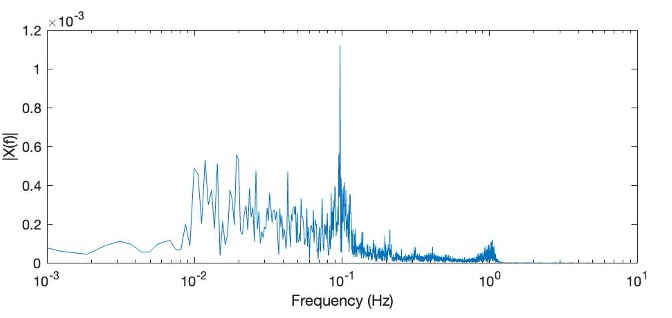

**Session 2:** 05_27

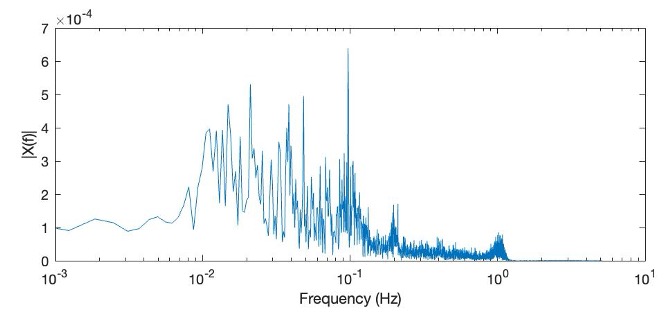

**Session 3:** 06_24

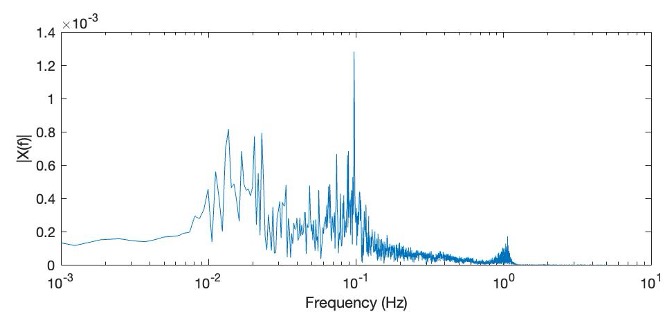

**Session 4:** 06_25

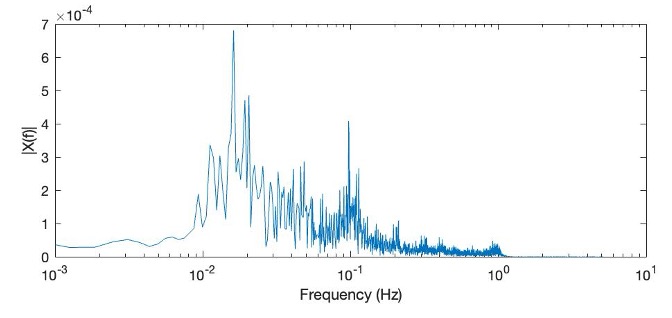

**Session 5:** 07_01

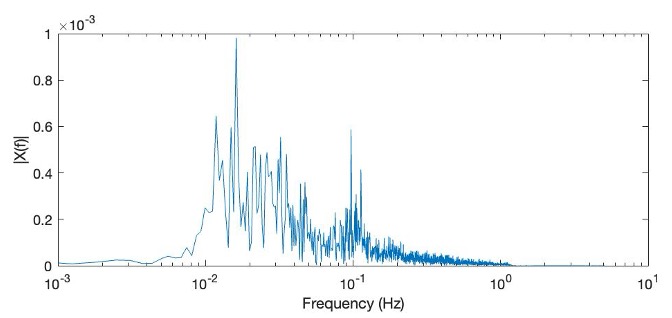

**F.** An additional low pass filter attenuates frequencies above 0.1 Hz to ensure the entirety of the heartrate is excluded from the stimuli. Note the drastically reduced high-frequency noise, leaving only waveforms at the stimulus frequency.

**Session 1:** 05_09

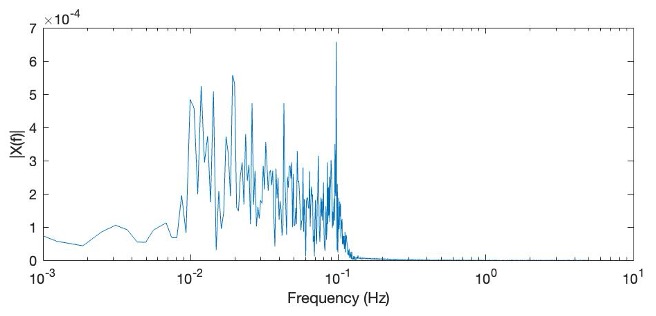

**Session 2:** 05_27

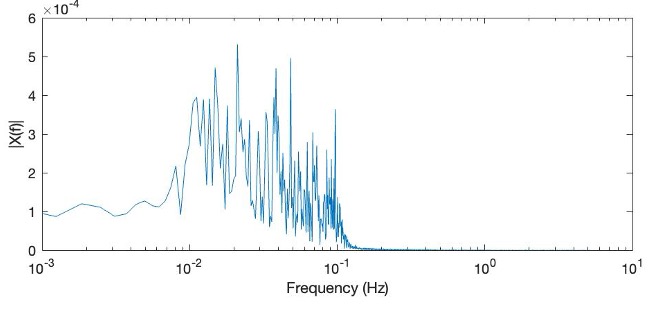

**Session 3:** 06_24

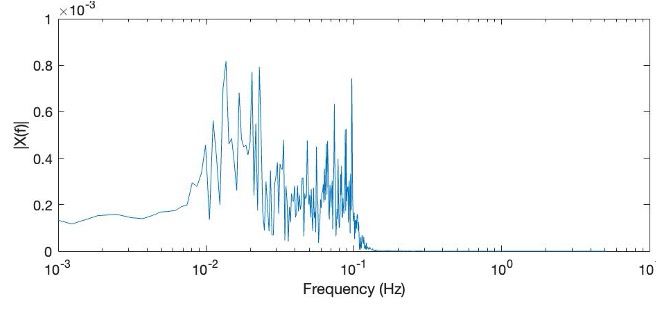

**Session 4:** 06_25

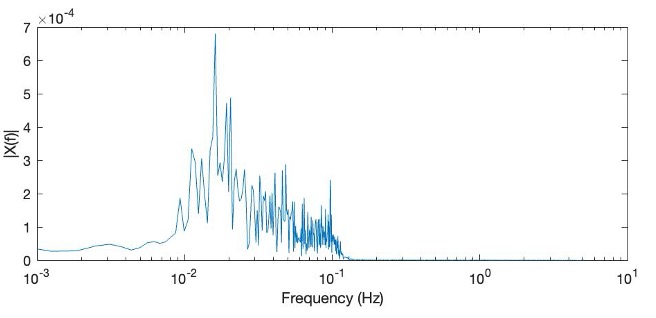

**Session 5:** 07_01

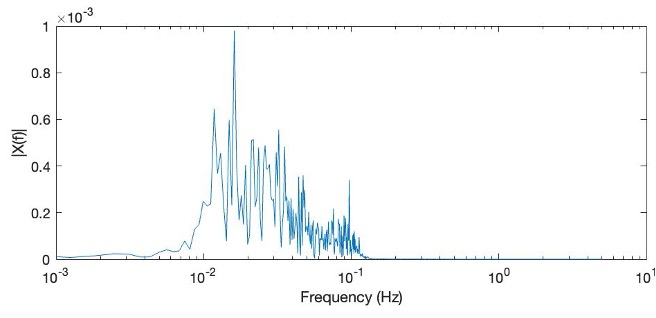

**G.** Finally the data from sessions one thorugh five are resampled at 1 Hz

**Session 1:** 05_09

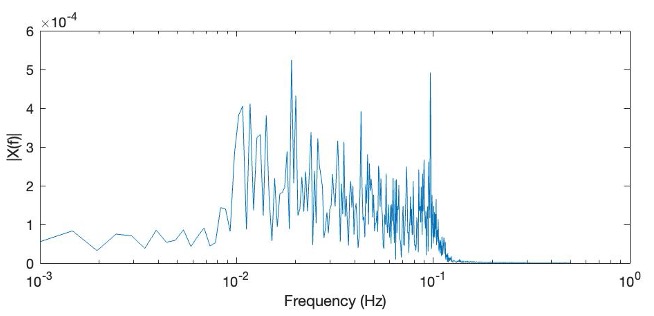

**Session 2:** 05_27

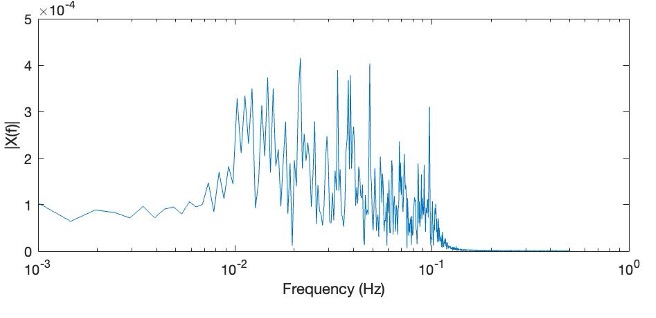

**Session 3:** 06_24

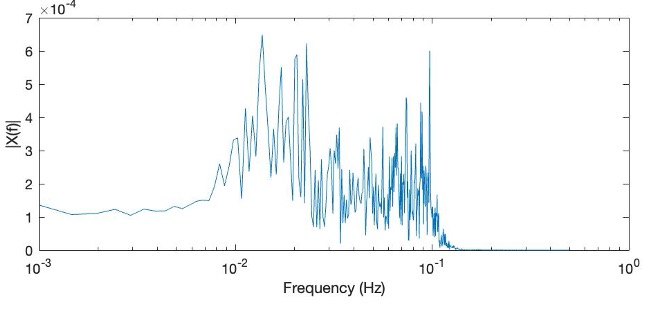

**Session 4:** 06_25

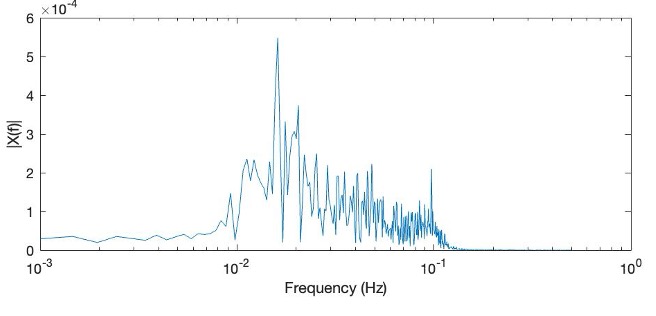

**Session 5:** 07_01

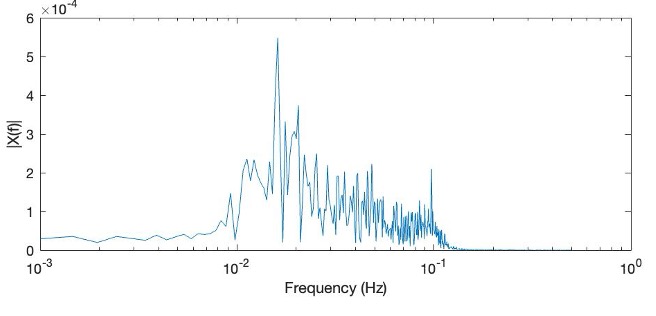
